# EdiProPred: Prediction of Edible Plant Tissue-Associated Proteins Using Large Language Models

**DOI:** 10.64898/2026.10.03.756411

**Authors:** Pushpendra Singh Gahlot, Akshay Shendre, Gajendra P. S. Raghava

## Abstract

Ensuring the safety and suitability of proteins considered for food and biotechnology applications is an important challenge. Computational approaches can facilitate the prioritisation of candidate proteins for further investigation, particularly given the rapid growth of genomic and metagenomic sequence datasets. In this study, we developed computational models to distinguish edible plant tissue-associated (EPTA) proteins from non-EPTA proteins. The positive class comprised 939 proteins with experimentally validated evidence of expression in plant tissues commonly consumed by humans, hereafter referred to as EPTA proteins. The negative class comprised 939 proteins annotated in UniProt as toxic, allergenic, or antinutritional, hereafter referred to as non-EPTA proteins. Models were evaluated using five-fold cross-validation with a cluster-based partitioning strategy, ensuring that no protein in the validation set shared more than 40% sequence identity with proteins in the corresponding training set. First, composition-based sequence features were evaluated to establish a baseline for EPTA or non-EPTA classification, with the Extra Trees (ET) classifier achieving a validation AUROC of 0.848. To incorporate evolutionary information beyond amino acid composition, PSSM profiles were subsequently evaluated, yielding an AUROC of 0.842 with an SVM classifier. Finally, to capture richer sequence patterns and contextual information using protein language modelling, we evaluated ESM2-based models. The fine-tuned ESM2-T33 model achieved the highest validation AUROC of 0.925 and was selected as the final predictive model. The resulting approach was implemented as the EdiProPred web server (https://webs.iiitd.edu.in/raghava/edipropred/) and standalone software for computational classification of plant proteins. The EdiProPred framework provides a useful computational tool for prioritising candidate plant proteins and facilitating the exploration of novel protein resources for food and biotechnology applications.

**Highlights:**

1. Proteins expressed in the edible parts of plants are considered edible proteins.
2. Toxins, allergens, and anti-nutritional proteins are considered non-edible proteins.
3. All models were trained, tested, and evaluated using non-redundant datasets.
4. The evolutionary language model achieved the highest performance.
5. A platform for the Prediction, Design, and Screening of Edible Proteins.

**Author’s Biography:**

1. Pushpendra Singh Gahlot is currently pursuing a PhD in Computational Biology at the Department of Computational Biology, Indraprastha Institute of Information Technology, New Delhi, India.
2. Akshay Shendre is currently working as a Research Assistant in the Computational Biology Department at the Department of Computational Biology, Indraprastha Institute of Information Technology, New Delhi, India.
3. Gajendra P.S. Raghava is currently working as a Professor and Director of Indraprastha Institute of Information Technology, New Delhi, India.

## 1. Introduction

Proteins are essential biomolecules that sustain life, serving as the primary source of amino acids for growth, repair, and metabolic functions (1). They are the building blocks for nearly every part of the human body, including muscles, skin, hair, and enzymes, and are vital for numerous physiological processes, such as immune response and tissue repair (2). Beyond their nutritional role, proteins also contribute to functional and bioactive properties that enhance food quality, human health, and therapeutic applications (3). With an estimated 8.6 billion people on the planet by 2050 and 11.2 billion by 2100, the sustainability of food systems around the world will face unprecedented challenges. Protein sources that are safe, sustainable, and nutritionally balanced are becoming more and more necessary in response to these growing demands and worries about food security (4). Although animal-based proteins are nutritionally important, their production puts a heavy burden on the environment, requiring large amounts of land and water and contributing significantly to greenhouse gas emissions (5). In this situation, plant-based proteins are becoming a great alternative, as they are more sustainable and can help meet nutritional needs while putting less pressure on the environment. Diets rich in plant proteins also tend to be lower in saturated fats and cholesterol, which have been linked to a lower risk of diseases like heart disease and type 2 diabetes (6).

Even though proteins are essential in our diet, not all of them are safe or suitable to eat. Some proteins can cause problems like allergies, toxicity, or may not be easily digested, which limits their use in food (7–9). Traditional protein sources like animal products also come with several challenges, including limited availability, environmental concerns, and ethical issues. These challenges have increased interest in alternative protein sources from plants, microorganisms, and engineered systems, creating a need for computational approaches that can facilitate the initial prioritisation and characterisation of candidate proteins for food-related applications.

To the best of our knowledge, no publicly available protein sequence database currently provides explicit labels indicating whether individual proteins are edible or non-edible, and no established sequence-based computational model is available for this classification task. Existing protein resources primarily provide annotations for specific properties, such as toxicity, allergenicity, nutritional characteristics, and biological function, rather than an integrated classification of proteins according to their association with edible plant tissues. Experimental assessment of these properties can also be costly, time-consuming, and resource-intensive. The absence of explicitly labelled sequence data and an established computational framework, therefore, represents a gap in the systematic computational prioritisation of plant proteins for potential food applications (10,11). This lack of robust predictive strategies creates a critical gap, hindering the effective utilisation of novel protein sources in food technology, biotechnology, and nutrition research.

To address this gap, we developed a comprehensive computational framework for distinguishing EPTA proteins from non-EPTA proteins (Figure 1). EPTA proteins were defined based on their reported occurrence in plant tissues commonly consumed by humans, whereas non-EPTA proteins were selected based on UniProt annotations for toxic, allergenic, or antinutritional properties. By integrating machine-learning approaches based on sequence-derived and evolutionary features with protein language models, we developed a computational strategy for EPTA/non-EPTA classification. The resulting framework provides a tool for the initial computational prioritisation of plant proteins for further investigation, with potential applications in food science, biotechnology, plant protein analysis, and sustainable nutrition.

**Figure 1:**
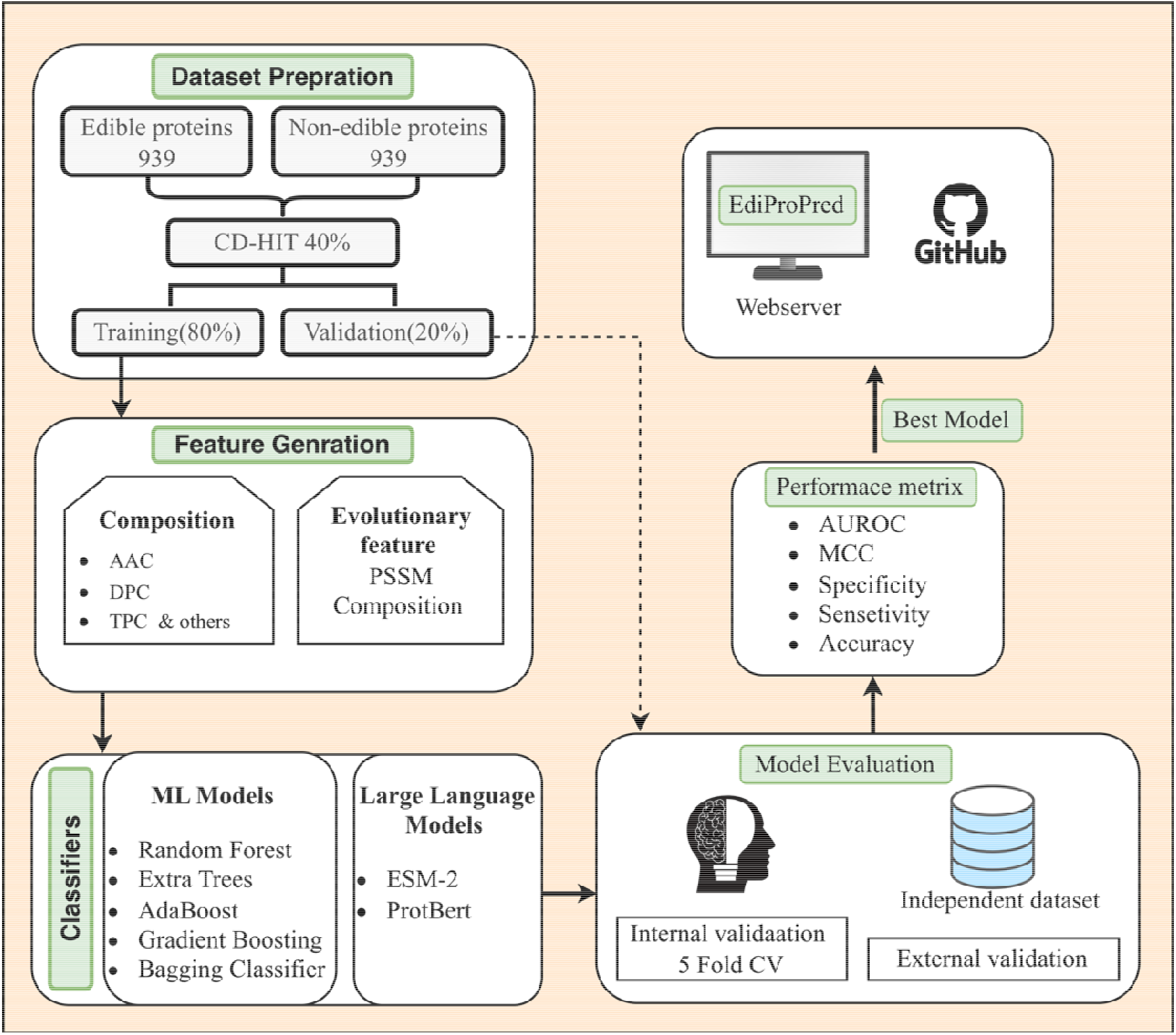
Flowchart illustrating EdiProPred’s entire computational pipeline

## 2. Materials and Methods

### 2.1 Dataset Formation

We obtained information on 35,103 plant species and their edible parts from the Food Plants International database (https://foodplantsinternational.com/). Proteins expressed in the edible parts of plants were then retrieved from UniProtKB/Swiss-Prot (12). Duplicate sequences and those shorter than 50 amino acids were excluded. To minimise redundancy, CD-HIT (13) was applied with a 90% sequence identity cutoff, yielding a final positive dataset of 939 proteins. For the negative dataset, harmful plant proteins annotated as toxic, allergenic, or antinutritional were retrieved from the UniProt database and restricted to the taxonomic group Viridiplantae (Taxonomy ID: 33090) as of March 10, 2025. The sequence length was restricted to 50-2,135 amino acids to match the length range of the positive dataset, and sequences outside this rang were excluded. The remaining sequences were clustered using CD-HIT at a 40% sequence identity threshold to remove redundancy, yielding 19,932 non-redundant proteins. From this set, 939 proteins were randomly sampled to match the size of the positive dataset. For the construction of Training and Validation Datasets, the positive dataset was first clustered using CD-HIT with a 40% sequence identity threshold, yielding 498 clusters. For the negative dataset, the 939 selected sequences were already non-redundant at 40% sequence identity, and therefore each negative sequence was treated as an independent cluster. The 498 positive clusters and 939 negative clusters were then distributed into five non-overlapping subsets using a round-robin strategy. In each fold, four subsets were combined to form the training dataset, while the remaining subset was used as the validation dataset. This cluster-based partitioning ensured that sequences in the validation dataset did not share more than 40% sequence identity with sequences in the corresponding training dataset, thereby minimising sequence redundancy and potential data leakage. The complete dataset formation and partitioning workflow is shown in Figure 2.

**Figure 2:**
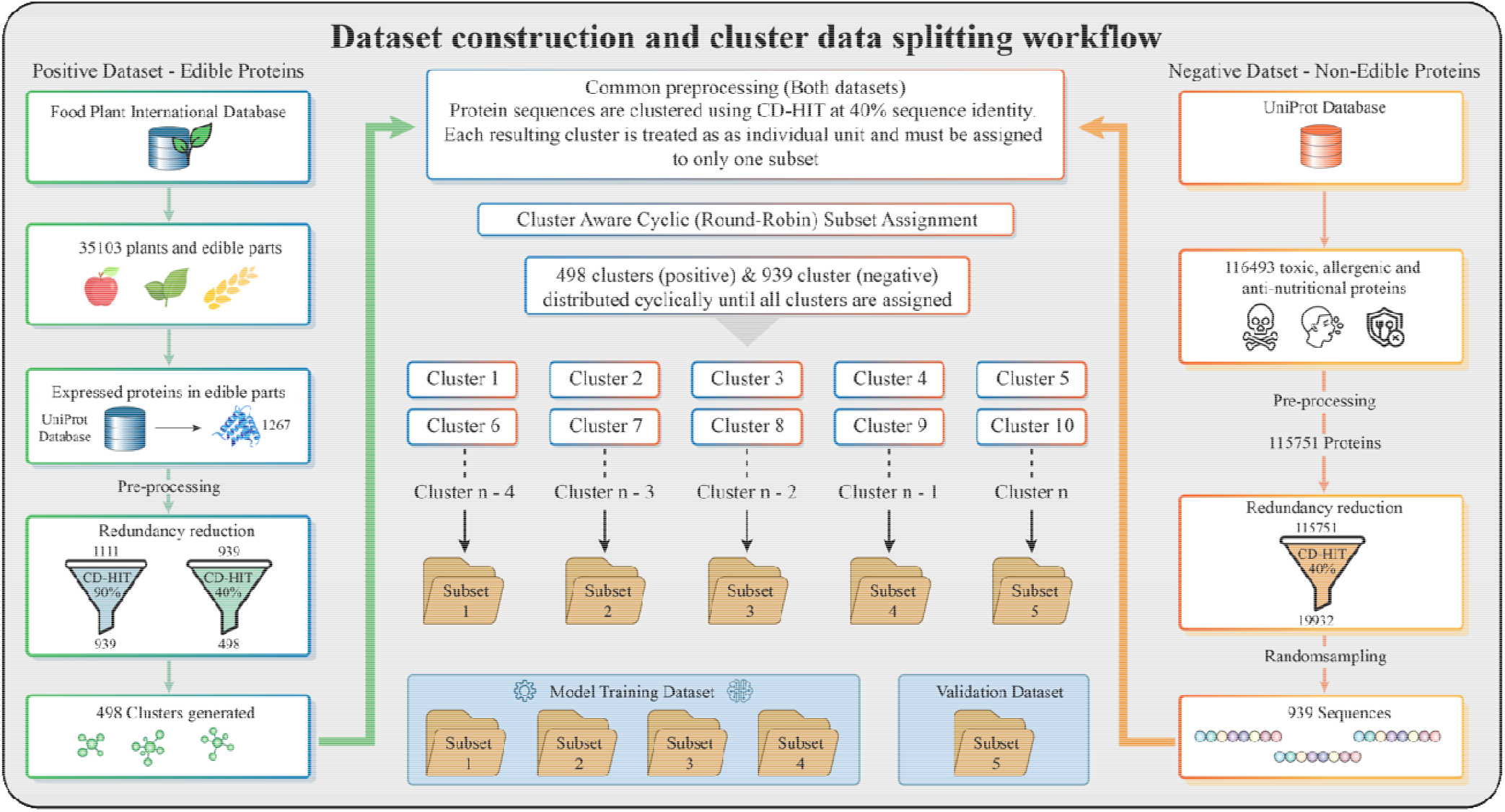
Overview of the EdiProPred Dataset Creation and Filtering Pipeline

### 2.2 Compositional Analysis

To better understand what differentiates EPTA and non-EPTA proteins, we carried out a detailed compositional and positional analysis. We used multiple analytical approaches to examine sequence-based properties across the datasets. In the first step, we focused on amino acid composition (AAC), where the percentage of each of the 20 standard amino acids was calculated for every sequence and then averaged separately for EPTA and non-EPTA proteins. To check whether the observed differences were statistically meaningful, we applied both the non-parametric Mann–Whitney U test and the parametric independent t-test. The resulting p-values were further adjusted using the Benjamini-Hochberg false discovery rate (FDR) method to correct for multiple comparisons. This approach helped us identify amino acids that show consistent and significant differences between the two groups.

### 2.3 Feature Generation

To develop a sequence-based prediction model, it is essential to effectively represent protein sequences in a way that captures their biological functions. We extracted a variety of informative features, including PSSM profiling, composition-based features, and word embeddings generated by large language models (LLMs), to ensure a comprehensive characterisation of the sequences.

#### 2.3.1 Evolutionary Information-based Feature

Leveraging evolutionary information to capture sequence conservation and variability, we characterised our sequences for their functional importance. Our feature extraction process began with the generation of a PSSM using Position-Specific Iterated BLAST (PSI-BLAST). The PSSM quantifies the evolutionary conservation of amino acid residues by comparing a sequence to its homologs. To obtain a uniform feature set, we then created a fixed-length vector of 400 elements, designated as PSSM-400, by summarising the compositional information of the PSSM profile.

#### 2.3.2 Composition-based Features

For composition-based feature extraction, we employed the standalone tool Pfeature, which generates a diverse range of sequence-derived descriptors. These included AAC, Dipeptide Composition (DPC), Atom Type Composition (ATC), Bond Type Composition (BTC), and a variety of physicochemical property (PCP) profiles, among others. Together, these descriptors capture both the global sequence makeup and underlying biochemical characteristics of proteins. A detailed summary of all extracted features, along with their vector dimensions, is provided in the Supplementary Table.

### 2.4 Machine Learning-based classifiers

After generating different feature representations, we applied several ML algorithms to classify proteins as EPTA or non-EPTA. We used Support Vector Machine (SVM) because it is effective in separating classes in high-dimensional data and can model both linear and non-linear relationships using kernel functions, making it well-suited for sequence-based problems. We also implemented Random Forest (RF), an ensemble method that builds multiple decision trees and combines their predictions to improve accuracy and reduce overfitting. In addition, ET, a variation of RF that introduces more randomness during tree construction, was evaluated to enhance generalisation. To capture more complex patterns in the data, we used Gradient Boosting (GB), which builds models sequentially by correcting the errors of previous ones. We also employed the K-Neighbours classifier (KN), which predicts a protein’s class based on the majority class among its nearest neighbours in the feature space. All models were implemented using the scikit-learn library, with parameters optimised through tuning, and their performance was evaluated using cross-validation and independent testing.

### 2.5 Large Language Models

Along with traditional ML methods, we also explored protein language models (PLMs) as direct classifiers. For this, we used ESM-2 and ProtBERT, two advanced transformer-based models that have been trained on large collections of protein sequences. These models treat protein sequences like a language, where amino acids act as tokens, allowing them to learn meaningful patterns and context within the sequence. By fine-tuning these models on our dataset, we adapted them to distinguish between EPTA and non-EPTA proteins. ProtBERT, which is based on the BERT architecture, was originally trained using a masked language modelling approach, helping it capture important structural and functional properties of proteins. Similarly, ESM-2, trained on UniRef protein datasets, has been shown to perform well across a variety of protein prediction tasks. Using these models allowed us to move beyond manual feature extraction and instead rely on rich representations learned directly from large-scale sequence data, ultimately improving prediction performance and generalizability.

### 2.6 Cross-Validation

To ensure the robustness and generalizability of our models, we employed 5-fold cross-validation, a widely adopted strategy in ML. In this approach, the dataset was randomly partitioned into k equal subsets (folds). For each iteration, one fold was used as the validation set, while the remaining k–1 folds were utilised for training. This process was repeated k times, ensuring that every sample in the dataset was used both for training and validation. The final performance of the model was obtained by averaging the results across all folds, which minimises the risk of overfitting and provides a more reliable estimate of predictive accuracy. The 5-fold cross-validation framework was particularly effective in minimising overfitting and ensuring that the results were not influenced by the bias of a single train-test split. This method also enabled us to fairly compare the performance of different classifiers and PLMs, thereby strengthening the reliability and reproducibility of our findings.

### 2.7 Performance Metrics Calculation

The ML models in this study were evaluated using a comprehensive set of performance metrics, covering both threshold-dependent and threshold-independent measures. The metrics considered include specificity, sensitivity, Matthews correlation coefficient (MCC), accuracy, and the area under the receiver operating characteristic curve (AUROC). Among these, AUROC is threshold-independent and provides an aggregate measure of the model’s discriminative ability across all classification thresholds. In contrast, specificity, sensitivity, MCC, and accuracy are threshold-dependent; therefore, their values were optimised by selecting the threshold that maximised performance.

These metrics were chosen as they provide complementary perspectives on model performance. Sensitivity and specificity highlight the model’s ability to correctly identify positive and negative samples, respectively, while MCC is regarded as a balanced metric even in cases of imbalanced datasets. Accuracy offers an overall performance measure, and AUROC ensures robustness by evaluating performance independently of the decision threshold. Collectively, these evaluation strategies ensure a reliable and unbiased assessment of the classification models, consistent with prior studies in the field. The following equations 1-4 were used to calculate these:

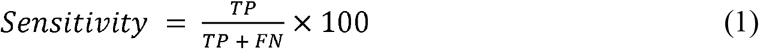

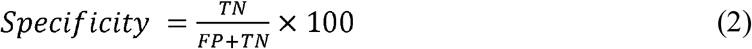

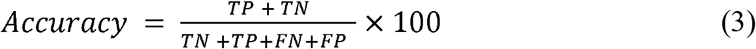

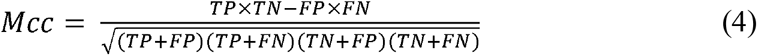

Where TP is a true positive, FP is a false positive, TN is a true negative, and FN is a false negative.

## 3. Results

### 3.1 Composition Analysis

We calculated AAC for EPTA and non-EPTA proteins, as shown in Figure 3. We observed that the amino acids Glutamic acid (E), Proline (P), and Alanine (A) are more abundant in EPTA proteins. Similarly, amino acids Serine (S), Asparagine (N), and Arginine (R) are widespread in non-EPTA proteins. We calculated AAC using the formula shown in Equation 5.

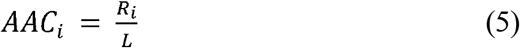

**Figure 3:**
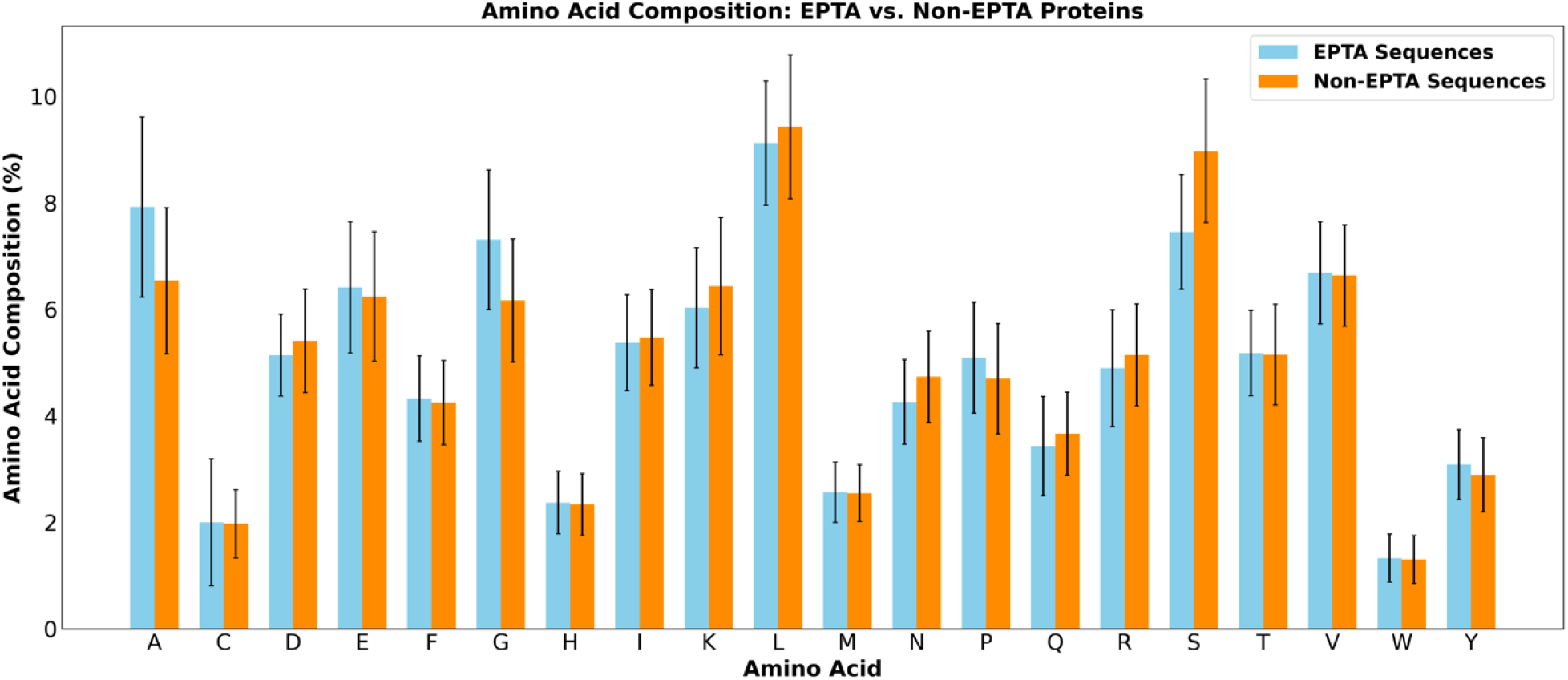
The percentage amino acid compositional analysis for EPTA or non-EPTA proteins.

### 3.2 Composition-Based Model

To investigate the role of sequence-derived information in EPTA protein prediction, we computed a wide range of compositional and order-based features using the pfeature package (14). Specifically, we extracted AAC, DPC, TPC, QSO, and other sequence descriptors. Each feature set was used independently to train ML classifiers, including SVM, RF, ET, GB, and KN. Among all tested features, the AAC descriptor consistently outperformed others, demonstrating its ability to effectively capture sequence-order information relevant for discriminating EPTA from non-EPTA proteins. The ET model trained on AAC features achieved AUROCs of 0.851 and 0.848 on the training and validation sets, respectively, underscoring its robustness and generalisation. Results based on AAC are summarised in Table 1, while performances of other feature sets (AAC, DPC, TPC, etc.) are provided in Supplementary Table S1. These findings highlight the advantage of incorporating compositional features, such as AAC, for reliable EPTA protein prediction.

**Table 1:** The performance of different machine learning-based models using the AAC feature of protein sequences.

| Models | Training Dataset |  |  |  |  | Validation Dataset |  |  |  |  |
| --- | --- | --- | --- | --- | --- | --- | --- | --- | --- | --- |
|  | Sen | Spec | Acc | AUROC | MCC | Sen | Spec | Acc | AUROC | MCC |
| **ET** | 0.804 | 0.762 | 0.783 | 0.851 | 0.566 | 0.768 | 0.770 | 0.769 | 0.848 | 0.538 |
| **RF** | 0.751 | 0.753 | 0.752 | 0.831 | 0.504 | 0.768 | 0.781 | 0.774 | 0.845 | 0.549 |
| **GB** | 0.759 | 0.723 | 0.741 | 0.823 | 0.482 | 0.796 | 0.701 | 0.747 | 0.818 | 0.498 |
| **SVM** | 0.778 | 0.718 | 0.748 | 0.820 | 0.497 | 0.829 | 0.701 | 0.764 | 0.837 | 0.533 |
| **KN** | 0.815 | 0.652 | 0.734 | 0.791 | 0.474 | 0.796 | 0.652 | 0.723 | 0.788 | 0.452 |

### 3.3 PSSM Feature-Based Model

To capture evolutionary information, we extracted PSSM profiles using the POSSUM (15) software and computed PSSM-composition features (PSSM-400), which encode the evolutionary propensity of each amino acid at each sequence position. These features were evaluated using RF, GB, SVM, LR, and ET classifiers, with the results summarised in Table 2. The SVM classifier trained on PSSM-400 features achieved AUROC values of 0.855 on the training set and 0.842 on the validation set. These results indicate that evolutionary information captured in PSSM profiles provides meaningful discrimination, and that PSSM-400 features, combined with an SVM classifier, provide a stable and reliable framework for distinguishing EPTA from non-EPTA proteins.

**Table 2:** The performance of different machine learning techniques-based models using the PSSM feature of protein sequences.

| **Models** | **Training Dataset** |  |  |  |  | **Validation Dataset** |  |  |  |  |
| --- | --- | --- | --- | --- | --- | --- | --- | --- | --- | --- |
|  | **Sen** | **Spec** | **Acc** | **AUROC** | **MCC** | **Sen** | **Spec** | **Acc** | **AUROC** | **MCC** |
| **RF** | 0.877 | 0.633 | 0.772 | 0.857 | 0.533 | 0.856 | 0.64 | 0.768 | 0.830 | 0.513 |
| **GB** | 0.836 | 0.697 | 0.777 | 0.852 | 0.541 | 0.818 | 0.672 | 0.758 | 0.818 | 0.495 |
| **SVM** | 0.841 | 0.716 | 0.787 | 0.855 | 0.564 | 0.895 | 0.664 | 0.801 | 0.842 | 0.583 |
| **ET** | 0.867 | 0.674 | 0.784 | 0.867 | 0.558 | 0.851 | 0.632 | 0.761 | 0.824 | 0.499 |
| **LR** | 0.833 | 0.696 | 0.774 | 0.843 | 0.536 | 0.851 | 0.656 | 0.771 | 0.828 | 0.520 |

### 3.4 Large Language Models

To further enhance prediction accuracy, we investigated the potential of Large Language Models (LLMs), which capture long-range dependencies and contextual information from protein sequences. Specifically, we fine-tuned two families of models: ESM2 (Evolutionary Scale Modelling) and ProtBERT. For ESM2, we evaluated multiple transformer depths (T6, T12, T30, and T33) to systematically analyse the effect of model complexity on predictive performance. Among these, the ESM2-T33 fine-tuned model achieved the best results with an AUROC of 0.925 on the validation set, outperforming smaller variants such as T6, T12, and T30. ProtBERT also demonstrated strong predictive power, although its performance was lower than ESM2-T33. These results highlight that fine-tuning large PLMs substantially improves the ability to distinguish EPTA from non-EPTA proteins compared to classical ML approaches using handcrafted features. The superior performance of ESM2-T33 suggests that deeper transformer architectures are particularly effective at capturing the nuanced sequence patterns that govern protein edibility. Detailed performance metrics are summarised in Table 3.

**Table 3:** The performance evaluation of PLMs on the validation dataset.

| PLMs | Sen | Spec | Acc | AUROC | MCC |
| --- | --- | --- | --- | --- | --- |
| **esm2_t33_650M_UR50D** | 0.862 | 0.866 | 0.864 | 0.925 | 0.728 |
| **esm2_t30_150M_UR50D** | 0.801 | 0.802 | 0.802 | 0.895 | 0.603 |
| **esm2_t12_35M_UR50D** | 0.796 | 0.791 | 0.793 | 0.889 | 0.587 |
| **esm2_t6_8M_UR50D** | 0.812 | 0.807 | 0.810 | 0.883 | 0.620 |
| **ProtBERT** | 0.801 | 0.802 | 0.802 | 0.899 | 0.603 |

### 3.5 Final Model

The final predictive model for EPTA protein identification was developed using the ESM2-T33-650M-UR50D protein language model, which achieved the highest validation performance, with an AUROC of 0.925. ESM2-T33 comprises 33 transformer layers and generates 1,280-dimensional residue-level representations. For each protein sequence, the representations from the final transformer layer were aggregated by mean pooling to obtain a fixed-length 1,280-dimensional sequence representation. These representations were subsequently passed to a fully connected linear classification layer with two output classes corresponding to EPTA and non-EPTA proteins. During training, the ESM2-T33 parameters were kept fixed, while the classification layer was optimised using cross-entropy loss and the Adam optimiser with a learning rate of 1 × 10^−4^. A batch size of 4 was used, with a maximum of 200 training epochs. Early stopping was applied based on the validation AUROC with a patience of 5 epochs, and the model with the highest validation AUROC was retained as the final model. For binary classification, a decision threshold of 0.55 was selected based on the threshold at which sensitivity and specificity were most closely balanced. The resulting model served as the final EPTA/non-EPTA classifier in EdiProPred.

## 4. Case Study: Large-Scale Prediction of EPTA Proteins in Poaceae Proteomes

To demonstrate the real-world applicability and scalability of EdiProPred, we conducted a comprehensive case study on the Poaceae family. A total of five complete Poaceae proteomes (*Oryza brachyantha, Setaria italica, Oryza sativa subsp. Indica, Hordeum vulgare subsp. Vulgare, Brachypodium distachyon, Sorghum bicolour*) were downloaded from the UniProt database and analysed using the standalone version of EdiProPred, powered by our best-performing model (fine-tuned ESM2-T33). In total, these proteomes contained 451,953 protein sequences. Each protein was classified as EPTA or non-EPTA based solely on its primary amino acid sequence. Of the full dataset, 196,764 proteins were predicted as potential EPTA proteins, while 255,189 proteins were identified as non-EPTA. This large-scale application demonstrates the robustness of the model and its ability to process complete proteomes efficiently and accurately. This case study highlights the practical utility of EdiProPred for high-throughput screening of plant proteomes. The tool can guide experimental prioritisation, support nutritional studies, and accelerate the discovery of safe and sustainable protein sources for future food systems.

## 5. Web Server Development of EPTA Protein Classification

To enable easy and widespread access to EPTA protein classification, a dedicated web server for EdiProPred was developed. The web server provides a simple and intuitive interface that allows users to submit protein sequences in FASTA or plain text format and obtain rapid predictions. Due to server-side computational limitations and the need to support multiple concurrent users with low-latency response times, the ESM2-T12 model was implemented for web-based inference. This model offers an effective balance between predictive performance and computational efficiency, making it well-suited for real-time EPTA protein classification in a web environment.

## 6. Discussion

Ensuring a continuous and sustainable supply of proteins is a global challenge, especially with the increasing population and rising demand for nutritious food. Traditionally, proteins have been obtained mainly from animal sources such as meat, eggs, and dairy products. However, animal-derived proteins face significant challenges, including environmental impacts, high production costs, limited scalability, and health concerns. Therefore, there is an urgent need to explore and utilise alternative protein sources that are safe, affordable, and environmentally sustainable. In this context, plant-derived edible proteins emerge as a promising solution. They not only provide balanced nutrition but also offer advantages in terms of sustainability, lower greenhouse gas emissions, and reduced resource consumption compared to conventional animal protein production. Moreover, certain bioactive and therapeutic proteins possess functional properties that may also be beneficial in dietary applications, provided they are safe and non-toxic((8). In addition, the concept of edible vaccines, where antigenic proteins are expressed in plants and consumed orally to induce an immune response, further highlights the potential of plant-based proteins in both nutrition and healthcare fields (16). Together, these developments emphasise the growing overlap between nutritional, therapeutic, and functional proteins, underscoring the importance of systematically identifying proteins that are both biologically active and suitable for human consumption (17,18).

The identification of edible versus non-edible proteins in plants is a key step toward expanding the range of dietary protein sources. Plant proteins can vary widely in digestibility, allergenicity, and nutritional profile, and therefore, distinguishing edible proteins from non-edible ones becomes critical. Experimental methods for such classification are time-consuming, costly, and often require large-scale validation studies. Computational approaches can play a transformative role by enabling rapid screening and prediction of protein edibility directly from sequence information. Our study contributes to this effort by developing EdiProPred, a computational framework for EPTA protein prediction that leverages a fine-tuned protein language model, ESM2-T33, to capture complex sequence representations for distinguishing EPTA from non-EPTA proteins.

We began with compositional analysis to identify amino acid preferences and distinguish EPTA from non-EPTA proteins. Notably, alanine (A) and serine (S) compositions showed significant differences between the two classes, reflecting their potential importance in determining digestibility and functional roles in proteins. Using sequence-based features such as AAC, DPC, and TPC, our models achieved encouraging performance, with the ET classifier trained on AAC features achieving AUROC scores of 0.851 (training) and 0.848 (validation). This confirmed that even simple composition-based features capture meaningful distinctions between EPTA and non-EPTA proteins. To further enhance prediction, we employed PSSM-based features derived from evolutionary information. Using POSSUM software, we generated PSSM profiles and extracted PSSM-400 composition features. Among multiple classifiers tested (RF, GB, SVM, LR, and ET), the SVM model achieved the highest AUROC values of 0.855 on the training set and 0.842 on the validation set. These results indicate that evolutionary conservation patterns captured through PSSM features support effective discrimination between EPTA and non-EPTA proteins. Moving beyond handcrafted features, we explored PLMs to leverage recent advances in deep learning. State-of-the-art models, including ESM2 with different transformer depths (T6, T12, T30, and T33) and ProtBERT, were directly fine-tuned on the EPTA versus non-EPTA protein classification task. Fine-tuning enabled the models to adapt their internal representations to the specific problem domain. Among the evaluated models, fine-tuned ESM2-T33 achieved the highest validation AUROC of 0.925, followed by ProtBERT. Based on a systematic comparison of model architectures, the fine-tuned ESM2-T33 model was selected as the final model due to its strong predictive performance.

While the ESM2-T33–based models form the core of the standalone tool, the web server employs the ESM2-T12 model, which is optimised for fast, server-side predictions. Together, these models constitute the backbone of EdiProPred, enabling flexible deployment across web-based and standalone environments and accommodating diverse computational requirements. The availability of EdiProPred as a web server and a standalone version ensures accessibility for a wide range of users, from computational biologists to experimental scientists. The web server provides a user-friendly interface for researchers without programming expertise, while the standalone version supports integration into custom computational pipelines. This multi-platform availability enhances the usability and applicability of the model in real-world scenarios. From a broader perspective, this study demonstrates that plant EPTA proteins can be systematically identified and classified using computational approaches, thereby accelerating the discovery of novel dietary protein sources. Such predictions can guide experimental validation, reduce unnecessary laboratory costs, and shorten the timeline for integrating new proteins into the food supply chain. Importantly, this work aligns with the global push toward sustainable agriculture and food systems, where plant-based proteins are expected to play a central role.

Despite these strengths, the present study has several limitations. The model is trained on currently available annotations, which may contain biases toward well-studied plant species and protein families. Additionally, edibility is influenced by complex biological and physiological factors, including post-translational modifications, processing methods, dosage, and host-specific responses, which are not fully captured by sequence-based models. While fine-tuned PLMs provide strong predictive performance, experimental validation remains essential to confirm the safety and nutritional suitability of predicted EPTA proteins. Future work will focus on expanding curated datasets, incorporating functional and experimental evidence, and improving model interpretability to further enhance the robustness and applicability of EdiProPred.

## 7. Conclusion

This study presents a comprehensive computational framework for distinguishing EPTA and non-EPTA plant proteins. By integrating traditional sequence-derived features, evolutionary information, and PLMs, we developed a robust predictive approach and deployed it as an accessible tool for the scientific community. In the context of increasing demand for sustainable food systems, such computational frameworks can support the identification and prioritisation of novel plant protein sources, thereby contributing to efforts toward global food security. Beyond this, the proposed approach has several important applications. It can assist in the rapid screening of plant proteomes to identify potential edible proteins, guide experimental validation and functional characterisation, and support the development of nutritionally enhanced food products. Additionally, it can be used in the discovery of bioactive and therapeutic proteins suitable for safe consumption, as well as in the design of edible vaccines and functional foods. The framework can also aid in risk assessment by filtering out potentially toxic or allergenic proteins, thereby improving food safety. Overall, EdiProPred provides a valuable computational resource for advancing research in food biotechnology, nutrigenomics, and protein engineering.

## Abbreviations

AAC: Amino acid composition
DPC: Dipeptide composition
TPC: Tripeptide composition
PSSM: Position-specific scoring matrix
RF: Random forest
ET: Extra Trees
KN: K-neighbours
GB: Gradient Boosting
SVM: Support vector machines
LR: Logistic Regression
Sen: Sensitivity
Spec: Specificity
Acc: Accuracy
AUROC: Area Under the Receiver Operating characteristic Curve
MCC: Matthews correlation coefficient

## Data availability statement

The dataset utilized in this study is publicly available through the EdiProPred web server (https://webs.iiitd.edu.in/raghava/edipropred/download.html). The corresponding source code can be accessed at the GitHub repository (https://github.com/raghavagps/EdiProPred).

## Author’s contribution

PSG and GPSR were responsible for dataset collection and processing. PSG developed the computational programs and implemented the algorithms and prediction models. PSG and AS jointly developed the front-end and back-end components of the web server. PSG, AS, and GPSR contributed to manuscript preparation. GPSR conceptualized the project, coordinated its execution, and provided overall supervision.

## Acknowledgments

The authors gratefully acknowledge the University Grants Commission (UGC) for providing fellowships and financial assistance, and the Indraprastha Institute of Information Technology Delhi (IIIT-Delhi) for providing the necessary infrastructure. The authors also acknowledge the Department of Biotechnology (DBT) for awarding an infrastructure grant to the institute. Adobe Illustrator and draw.io were used to prepare the figures presented in this work. Additionally, the authors acknowledge the use of ChatGPT (OpenAI) for assistance with language editing of the manuscript.

